# Crystallographic and NMR studies of Streptococcus pneumonia LCP protein Psr_Sp_ indicate the importance of dynamics in four long loops for ligand specificity

**DOI:** 10.1101/2024.10.21.619401

**Authors:** Tatyana Sandalova, Benedetta Maria Sala, Martin Moche, Hans-Gustaf Ljunggren, Evren Alici, Birgitta Henriques-Normark, Tatiana Agback, Dmitry Lesovoy, Peter Agback, Adnane Achour

## Abstract

The crystal structure of the extracellular region of the second pneumococcal LCP, a polyisoprenyl-teichoic acid-peptidoglycan teichoic acid transferase Psr_Sp_, was determined and refined to 2.15Å resolution. Despite the low sequence homology with other LCP proteins, the Psr_Sp_ maintains the fold of the LCP domain and the positions of the 15 residues suggested to participate in the transferase function are conserved. The empty tunnel found in the Psr_Sp_ between the central β-sheet and three α-helices is wide enough to accommodate polyisoprenyl-teichoic acid. Comparison of the crystallographic temperature factors of LCP from distinct bacteria demonstrated that the four long loops located close to the teichoic acid and peptidoglycan binding sites have different relative mobility. To compare the dynamics of the Psr_Sp_ in crystalline state and in solution, NMR spectra were recorded, and 88% of the residues were assigned in the ^1^H-^15^N TROSY HSQC spectra. Comparison of the secondary structure of the crystal structure of Psr_Sp_ with NMR data demonstrated a perfect concordance between the results using these two methods. Moreover, the relative mobility of the essential loops estimated from the crystallographic B-factor is in good agreement with order parameter S^2^, predicted from chemical shift. We hypothesize that the dynamics of these loops are important for the substrate promiscuity of LCP proteins.

## Introduction

Glycopolymers in Gram-positive bacteria play a crucial role in the maintenance of structural integrity and functionality of the cell wall, significantly contributing to the bacterial ability to interact with their environment and to resist external stresses (1–3). In addition to the defense of the cell against the immune system of the host, bacterial glycopolymers bind to a large ensemble of other cell surface proteins, regulating their functional activity. They also participate in transport processes, cation homeostasis, resistance to antimicrobial peptides and lysozymes, regulation of cell elongation and cell division, as well as many other additional processes (4). Although the composition of these glycopolymers is very different in different bacteria, there are some common important features in their structure and their biosynthesis (3).

Teichoic acids (TA) are important cell wall polymers present in almost all Gram-positive bacteria. TAs are commonly divided into two classes; (i) lipoteichoic acid (LTA), which are anchored in the cytoplasmic membrane using a lipid anchor, and (ii) wall TA (WTA) that are covalently bound to the peptidoglycan (PG). Importantly, TA in *Streptococcus pneumoniae* is a major cell wall polymer with a unique identity, including several unusual sequence and structural features compared to other Gram-positive bacteria. First, the chemical composition of both LTA and WTA is identical in *S. pneumoniae*. Secondly, the *S. pneumoniae* TA is unusually complex and contains the rare positively charged amino sugar 2-acetamido-4-amino-2,4,6-trideoxy-D–galactose (AATGal) (5). Third, pneumococcal TA contains phosphorylcholine attached to the N-acetyl-galactose-amine residues, which is a binding site for several choline-binding proteins (6). Altogether, pneumococcal TA comprise four to eight repeating units (RU), all composed of AATGal, D-glucose, ribitol-phosphate, and two N-acetyl-galactose-amine residues modified by phosphorylcholine (7).

The biosynthesis of streptococcal TA RUs and their polymerization takes place on the cytoplasmic side of the bacterial cell membrane, and involves at least 18 different cytosolic and membrane-associated enzymes (7). The polymerization of TA-RUs occurs on the membrane-bound C55-undecaprenyl lipid carrier. When the synthesis of these RUs is completed, the transmembrane transporter TacF presumably moves the C55-PP-linked TA chains across the membrane. The last step of the WTA synthesis is implemented by a specific set of transferases from the LytR-CpsA-Psr (LCP) enzyme family which catalyzes the transfer of TA from the lipid carrier to the PG. Most often, covalent linkage occurs via the formation of a phosphodiester bond to the C6-OH group of the N-acetylmuramic acid of the PG (8). LCP transferases can catalyze not only TA attachment to the PG in *S. pneumoniae* but are also responsible for the linkage of many capsule polysaccharides to the PG, which plays a crucial role in the bacteria structure and pathogenicity (5, 9).

The LCP proteins are transmembrane proteins with a catalytically active extracellular C-terminus and cytoplasmic N-terminus. They have been described as important potential antimicrobial targets for Gram-positive pathogens due to the importance of their multiple functions, as well as their conserved structures (9). Several independent studies have revealed that a large array of different Gram-positive bacteria commonly have multiple copies of LCP enzymes that are seemingly functionally redundant. Three different genes encoding for LCP proteins have been hitherto identified in *Streptococcus pneumoniae, Streptococcus agalactiae, Staphylococcus aureus* and *Bacillus subtilis* (9, 10). It was shown for *B. subtilis* that the deletion of a single LCP member had no or only a minor effect on cell growth under normal conditions. However, removal of all three LCP proteins is lethal to the bacteria (10). In contrast, it should be noted that the deletion of even a single LCP gene in *S. pneumoniae* or *S. agalactiae* results in a reduced capsule volume, significantly decreased viability over time and reduced cell wall integrity (9, 11).

The three-dimensional structures of several LCP molecules from a large ensemble of bacterial species have been previously determined, including LcpA from *S. aureus* (6eux.pdb) (12), Cps2A from *S. pneumoniae* (2xxq.pdb, 4de8.pdb, 3tep.pdb) (9, 10), *S. agalactiae* (3okz.pdb), TagT (4de9.pdb, 6mps.pdb) (9, 13), TagV (6uf3.pdb) and TagU (6uf6.pdb) all from *B. subtilis* (12). Although the sequence identity among LCPs is low, ranging between 20-40% (**Figure 1**), the overall fold of the determined LCP domains is similar. The core of each determined LCP molecule has an α-β-α fold that contains a central β-sheet surrounded on both sides by several α-helices (**Figure 2**). Importantly, since the active site of different LCP proteins is surrounded by several flexible loops with different sequences, an enhanced understanding of the dynamic behavior of the active site is in our opinion crucial for adequate drug discovery targeting LCP proteins and/or a specific LCP protein.

**Figure 1.**
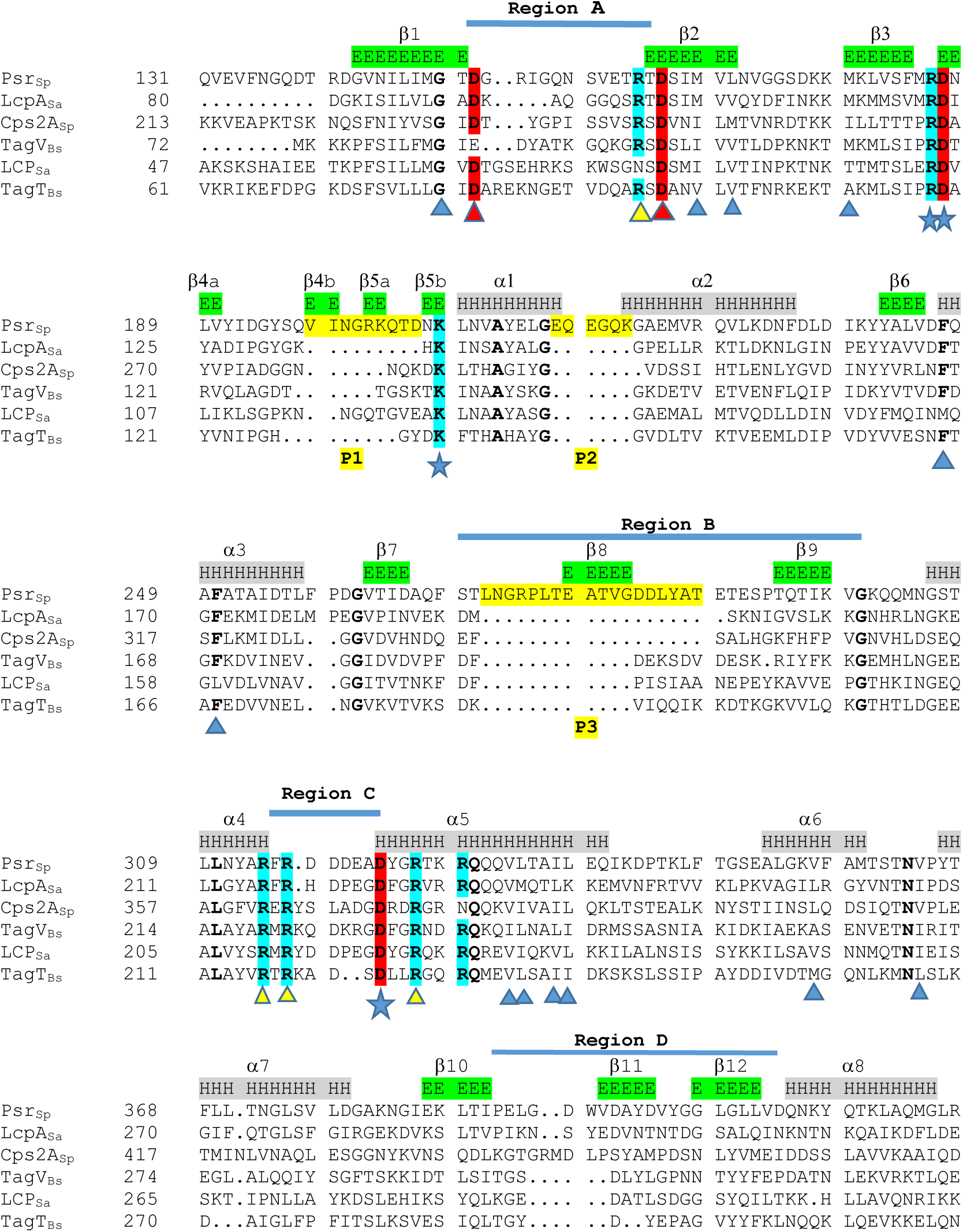
Multiple sequence alignment of the LCP domain of Psr_Sp_ with the five best structural hits found by DALI demonstrates low sequence conservation. A DALI search based on the crystal structure of Psr_Sp_ gave LcpA_Sa_ from *Staphylococcus aureus* (6uex.pdb), the LCP domain of Cps2A from *S. pneumoniae* (4de8.pdb), TagV from *B. subtilis* (6uf5.pdb), TagT from *B. subtilis* (6uf5.pdb), and the LCP protein from *S. agalactiae* (3okz.pdb) as the five best hits. Sequence alignment of these six proteins reveals low sequence conservation. The secondary structure of Psr_Sp_ and the localization of the A, B, C, D regions are presented above the sequences. The P1-P3 insertions in Psr_Sp_ are marked in yellow. The 16 residues that are strictly conserved among all six structures are in bold. The essential acidic residues within the active site are in red, while basic residues are in blue. The aspartic acid residues involved in Mg^2+^ binding are marked with red triangles, while the arginine residues that are essential for pyrophosphate binding are marked with yellow triangles. Residues lining the tunnel for the polyprenyl moiety of the substrate are marked with blue triangles. Residues that can be essential for peptidoglycan binding are marked with stars.

**Figure 2.**
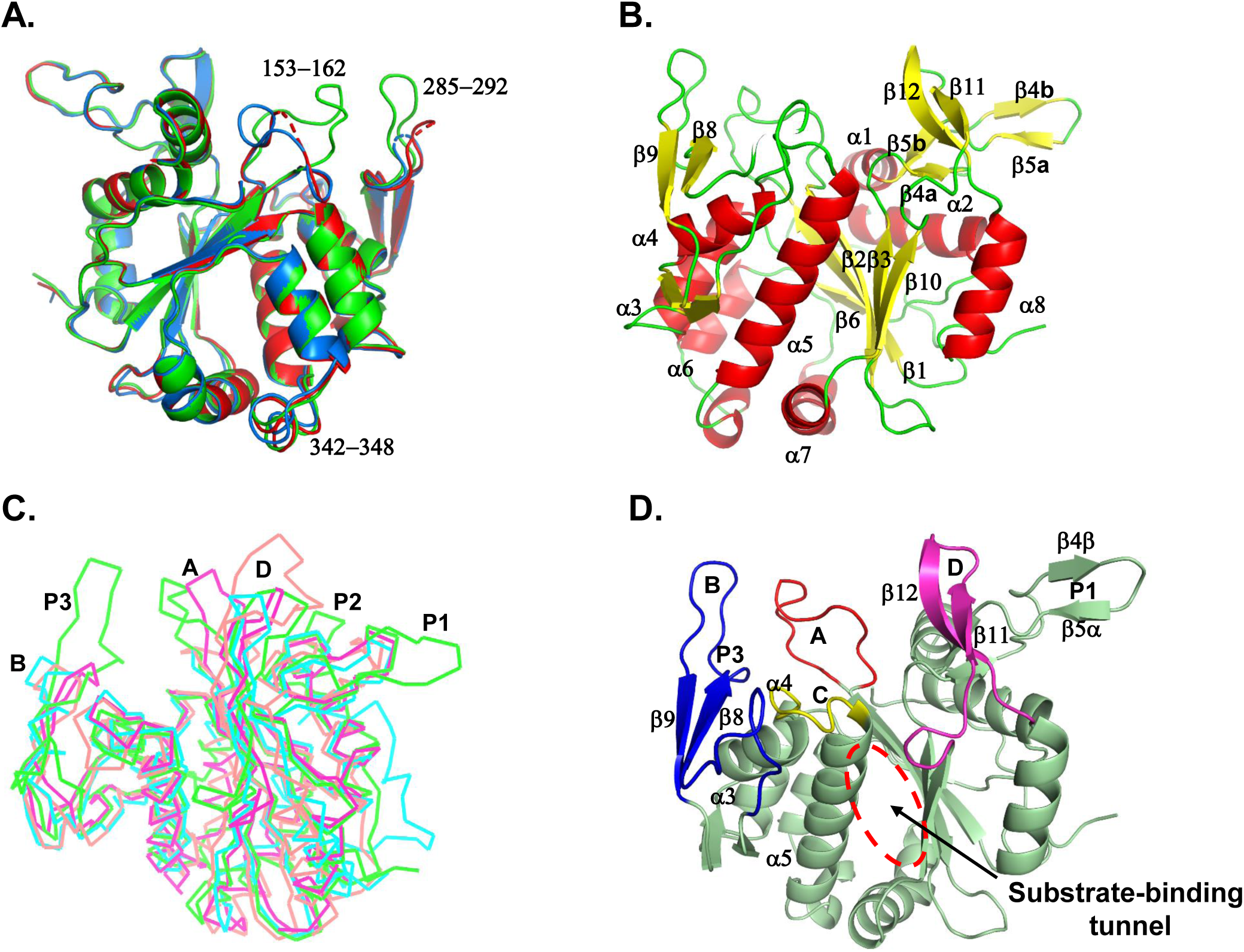
Crystal structure of the *S. pneumoniae*-associated Psr_Sp_. **A.** Superposition of the three Psr_Sp_ molecules found in the asymmetric unit of the crystal structure indicates flexibility in three different regions annotated by the corresponding residues. The three Psr_Sp_ molecules A, B and C found in the assymetric unit are in red, blue and green, respectively. **B**. The crystal structure of Psr_Sp_ reveals a typical overall LCP fold despite low sequence homology to other members of the LCP protein family. **C**. Superposition of Psr_Sp_ (in green) on the crystal structures of the three most similar LCP, LcpA_Sa_ from *Staphylococcus aureus* (in blue, 6uex.pdb), TagT from *Bacillus subtilis* (in red, 6uf5.pdb) and Cps2A from *S. pneumoniae* (in orange, 4de8.pdb) demonstrates that the main difference between these LCP molecules resides in the length of the P1-P3 protrusions and in the conformations of four different regions, that we named A, B, C and D. **D**. The A, B, C and D regions of Psr_Sp_, suggested to be important for substrate binding, are displayed in red, blue, yellow and magenta, respectively. The localization of the substrate binding tunnel is indicated.

The crystal structure of the *S. pneumoniae* LCP protein Cps2A has been previously determined, revealing how it binds to polyisoprenyl-pyrophosphate, a crucial step for attaching capsular polysaccharides to the cell wall PG (10). However, although the 3D-structure of Cps2A provided fundamental insights into how this enzyme catalyzes the final steps of the polysaccharide attachment process, the molecular bases underlying the importance of each *S. pneumoniae* LCP for attaching TA on either the bacterial cell wall or PG remain unclear. Here, in an effort to enhance our understanding of these non-excluding and possibly redundant functions, we determined the crystal structure of the extracellular region of the second pneumococcal LCP, the polyisoprenyl-teichoic acid-peptidoglycan teichoic acid transferase Psr_Sp_, (previously named MsrM), combining static high-resolution X-ray structural information with dynamic NMR analyses. We report the near-complete assignment of the ^1^H, ^13^C, and ^15^N backbone resonances for the Psr_Sp_ domain, as well as the ^13^Cβ side chain assignment of this protein. To resolve ambiguities in the crowded areas of the Psr_Sp_ LCP domain spectra, we compared the assigned resonances with those at different stages of deuterium exchange in the amide groups. Based on this data, the secondary structure of the Psr_Sp_ in solution was determined and compared with the crystal structure of the same protein, allowing us to evaluate the flexibility of this protein. Our results provide a structural basis for understanding the intra- and intermolecular interactions of the LCP domain of Psr_Sp_.

## Results

### The crystal structure of Psr_Sp_ reveals a classical LCP fold with the presence of three protruding regions

The *S. pneumoniae* Psr_Sp_ has a short N-terminal cytoplasmic region (residues 1-90), a single transmembrane helix (comprising residues 91-114) and an extracellular catalytic LCP domain encompassing residues 115-428. The construct used within the present X-ray crystallography study of Psr_Sp_, comprises almost the entire extracellular region of this protein (residues 130-424) and a C-terminal His-tag (**Figure 1**). Size exclusion and circular dichroism analyses demonstrated that Psr_Sp_ is a monomer with a molecular weight of 33 kDa and significant overall stability, with a melting temperature T_m_ of about 63°C. The crystal of Psr_Sp_ belongs to the space group P321 (**Table S1**), and contains according to solvent content analysis (14) three Psr_Sp_ molecules in the asymmetric unit. The crystal structure of Psr_Sp_ was determined by molecular replacement. Interestingly, the program Phaser (15) was unable to find any solution with initial models from any of the previously determined crystal structures of LCP domains, including Cps2A, the first structurally determined member of the pneumococcal LCP (2xxq, (10) nor LcpA from *S. aureus* (6eux, (12)). However, two molecules of Psr_Sp_ were unambiguously identified by Phaser using a molecular model obtained from the AlphaFold server (16). The third molecule in the asymmetric unit was found only upon using MolRep (17), after establishing the first two solutions with Phaser. The final crystal structure of Psr_Sp_ was refined to 2.15 Å resolution.

Although the three Psr_Sp_ molecules found in the asymmetric unit display highly similar three-dimensional structures, specific differences were identified in their backbones, testifying to the flexibility of this protein. Superposition of molecules A and B on C resulted in root mean square deviation (rmsd) values of 0.5 and 0.49 Å, respectively. The main difference between the three molecules relates to the conformation of the loop 153-162 which is similar in molecules A and B but very different in molecule C (**Figure 2A**). Furthermore, there is a gap corresponding to the stretch of residues 288-291 in the final model of molecules A and B, though this region is well defined in molecule C (indicated by dashed red and blue lines, respectively, in **Figure 2A**). In addition, the backbone conformation of residues 342-348 is different in molecule B compared to molecules A and C, due to close contacts with symmetry-related molecules in the crystal asymmetric unit. Since the final model of subunit C comprises all 298 amino acids of our construct, we decided that the description of the crystal structure of Psr_Sp_ will be performed using this chain. All three subunits form crystallographic trimers around the three-fold axis of P_321_. The 700 Å^2^ interface between neighboring subunits is formed by the N-terminal stretches of residues 130-139 and 368-390. However, we believe that this hypothetical trimer formation mainly reflects a crystallographic artifact since Psr_Sp_ behaves as a monomer during purification. NMR provides an alternative way, compared to gel filtration analyses, to discriminate monomeric and multimeric protein forms in solution, through the direct estimation of the rotational correlation time *τ*_C_. The experimental *τ*_C_ can be compared with the values predicted using an empiric equation that takes into account the temperature and molecular weight (18). Such estimates at 35°C gave *τ*_C_ values of 15.6, 31.1 and 46.6ns at for the monomeric, dimeric and trimeric forms of Psr_Sp_, respectively. The former value agrees well with 14.7ns obtained at 35°C by us using tract type NMR expert (19), thus confirming the predominantly monomeric form of Psr_Sp_ in solution.

The central core of the extracellular LCP domain of Psr_Sp_ is composed of a five-stranded β-sheet (β6-β1-β2-β3-β10) with the middle β2-strand running antiparallel to all other β-strands (**Figure 2B**). The central β-sheet is surrounded by eight α-helices, three of which, α1, α2, and α8, are located on one side of the β-sheet, and five additional α-helices on the other side. Analysis of the crystal structure of Psr_Sp_ using the Dali server (20) demonstrated the similarity of its fold to other previously determined LCP domain crystal structures. The structures that are most similar to Psr_Sp_ are the *S. aureus*-associated LcpA_Sa_ (6uex.pdb) (12), TagT from *B. subtilis* (6mps.pdb) (13) and Cps2A from *S. pneumoniae* (4de8.pdb) (9). The cores of these three proteins consisting of 230 residues align well to Psr_Sp_, with rmsd values of 2.0 Å for LcpA_Sa_, 2.7 Å for TagT_Bs_ and 2.3 Å for Cps2A. Superposition of these four proteins clearly demonstrates that despite their similarity in the overall fold, the sequence identity among LCP is not high, and pairwise alignment shows only 37% of identical residues between Psr_Sp_ and LcpA_Sa_, and 21-22% with TagT and Cps2A. Multiple sequence alignment of the seven proteins with the most similar three-dimensional structures to Psr_Sp_ confirms the overall low sequence homology among these members of the LCP family, with only 14 residues that are absolutely conserved (**Figure 1**). Superposition of the crystal structure of Psr_Sp_ with three other LCP proteins illustrates that there are three β-hairpin-like protrusions in Psr_Sp_, which are not present in other LCP proteins (**Figures 1 and 2D**). Protrusion P1 composed of the stretch of residues 188-206, with two antiparallel β-strands β4a-β5b and β4b-β5a, separates the β3-strand from the helix α1. Despite the insertion of nine residues, the conserved essential lysine residue K208 superposed very well with lysine residues on other LCP proteins (**Figure 1**). The relatively short protrusion P2 (residues 217-222) is located between helices α1 and α2. Finally, the longest protrusion P3 comprises residues 271-288 (**Figure 2C**). It should be noted that the length of these protruding chains is also significantly different among these LCP proteins.

### The substrate-binding site of Psr_Sp_ comprises common LCP features as well as highly unique properties

A wide and long empty tunnel is present in Psr_Sp_ between the central β-sheet and helices α3, α4 and α5 (**Figure 2D**). It should be noted that unlike LcpA_Sa_ and TagT, Psr_Sp_ was expressed and crystallized without any endogenous hydrophobic undecaprenyl molecule. However, the compound C_40_-PP-GlcNAc found in LcpA_Sa_ can be easily modelled inside the tunnel of Psr_Sp_ by simple superposition of Psr_Sp_ onto LcpA_Sa_. Small changes in the conformation of only three residues, D165, R314 and L337, are enough to remove all steric clashes for the appropriate binding of the ligand C_40_-PP-GlcNAc docked to substrate-free Psr_Sp_ (**Figure 3**). While the side chains of residues D165 and R314 localized in the binding site have to move to allow adequate binding to the polar moiety of the substrate binding, the conformation of the side chain of residue L337 positioned on helix α5 has to be changed to allow binding to the lipid part of C_40_-PP-GlcNAc.

**Figure 3.**
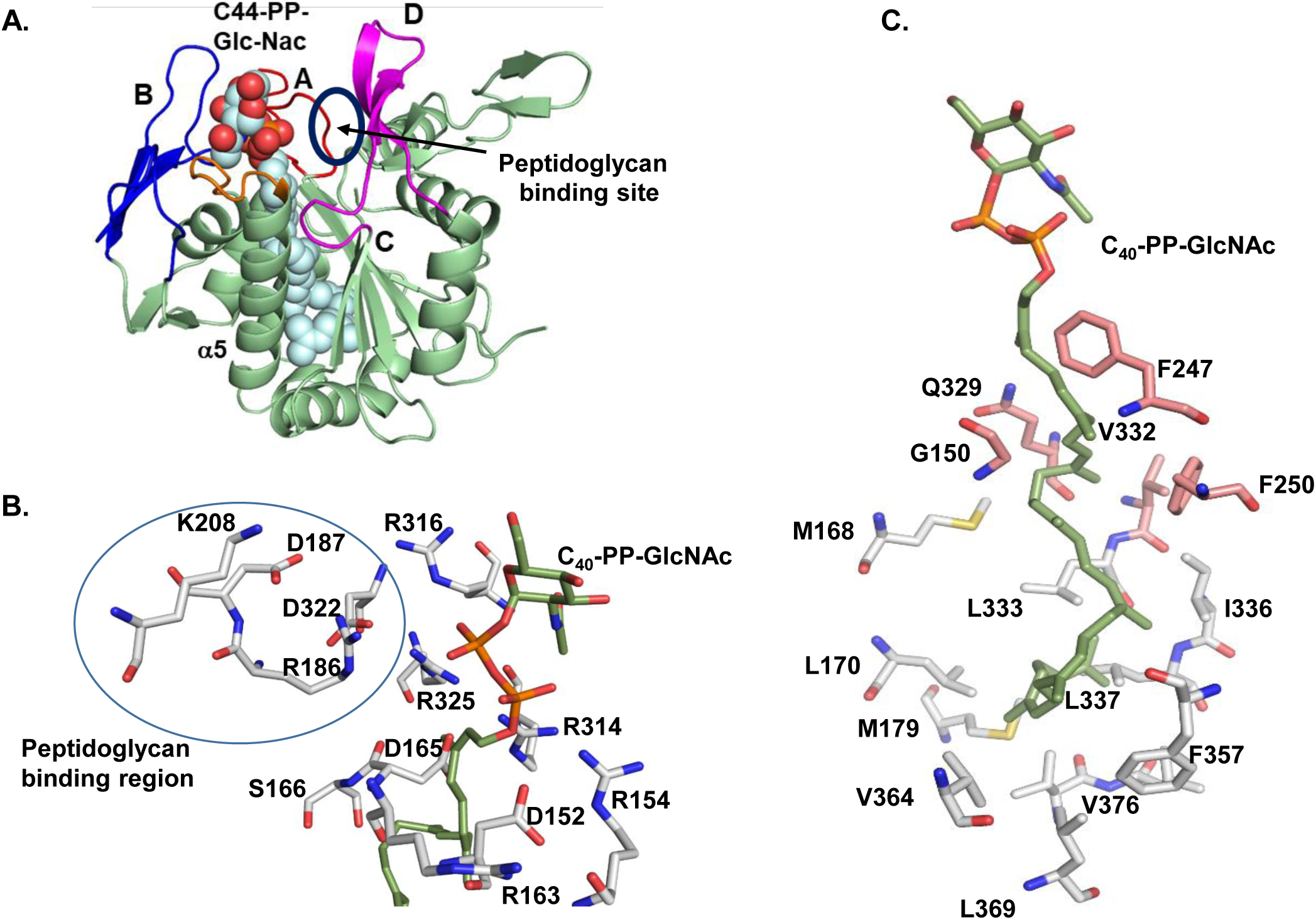
Superposition of LcpA_Sa_/C_40_-PP-GlcNAc onto ligand-free Psr_Sp_ reveals that the ligand fits well to the substrate-binding site of Psr_Sp_ and that all catalytic residues are located appropriately. **A.** The C_40_-PP-GlcNAc ligand found in the active site of LcpA_Sa_, is displayed as light blue spheres inside the tunnel within Psr_Sp_. The four regions A-D that are suggested to be important for substrate binding are colored in red, blue, orange and magenta, respectively. **B.** LCP-conserved residues are located in Psr_Sp_ close to the pyrophosphate moiety of the ligand. **C.** The hydrophobic part of the ligand is surrounded either by conserved residues (pink) or non-conserved but still hydrophobic residues (grey) of Psr_Sp_.

A detailed description of the substrate-binding site of LcpA_Sa_ and probable mechanisms underlying the function of LCP transferases has been previously provided (10, 12). The present crystal structure demonstrates that the catalytic mechanism of Psr_Sp_ is most probably highly similar to other LCP proteins. Indeed, despite the low sequence identity within LCP proteins, all twelve amino acids that have been suggested as important for the teichoic acid transferase activity are conserved in Psr_Sp_ (**Figures 1, 3B and 3C**). Though the residues that bind the pyrophosphate moiety of the ligand are highly conserved in LCP, the four loop-like regions that surround the active site are very different. Following previous descriptions (12), we will call these regions A (residues 152-164), B (267–298), C (315–322), and D (390–410) (**Figures 1 and 2D**). The B region of Psr_Sp_ includes the protrusion P3 which makes region B the longest among LPC proteins (**Figures 2C and 2D**). Regions A and C form loops that connect strands β1 and β2, and helices α4 and α5, respectively. Multiple sequence alignment (**Figure 1**) and superposition of 3D structures (**Figure 2C**) demonstrate that the length and conformation of all these four regions are different among LCP proteins. Several residues suggested to be essential for the catalysis activity, such as D152, R163, D165 and S166, are located on the N- and C-termini of region A. The shorter region C is also located between the four essential residues R314, R316 from helix α4, and D322, R325 from helix α5. Regions B and D are the most divergent among LCPs, since they do not comprise any conserved residue, which can reflect the different chemical structure of TA from different bacterial species. Calculation of electrostatic surface potentials demonstrates that the entrance to the tunnel is positively charged in all LCP molecules (**Figure 4**). A molecular model of the Psr_Sp_/C_40_-PP-GlcNac complex demonstrates that the five positively charged residues, R154, R186, R314, R316, and R325 are located at the entrance of the active side, ready to interact with the PP moiety of the TA-PP-polyisoprenyl molecule (**Figure 3B**). Comparison of the electrostatic potential on the surface of other LCP proteins emphasizes also the low conservation of surface residues outside the entrance (**Figure 4**). Psr_Sp_ is the most acidic within this region compared to LcpA_Sa_, TagT or Cps2A. This is not surprising since the overall number of acidic residues in the LCP domain of Psr_Sp_ amounts to 41, with only 30 basic residues, which stands in sharp contrast to LcpA_Sa_ which has 30 acidic and 37 basic residues. The main contribution to the overall acidity of the Psr_Sp_ surface is provided by regions B, C, and D which contain five, five and four negatively charged non-conserved residues, respectively. Since the WTA of *S. pneumoniae* contains the positively charged sugar AATGal which must be connected to the N-acetyl muramic acid (NAM) of PG, the presence of these acidic residues close to the active site is favorable for binding of WTA to Psr_Sp_.

**Figure 4.**
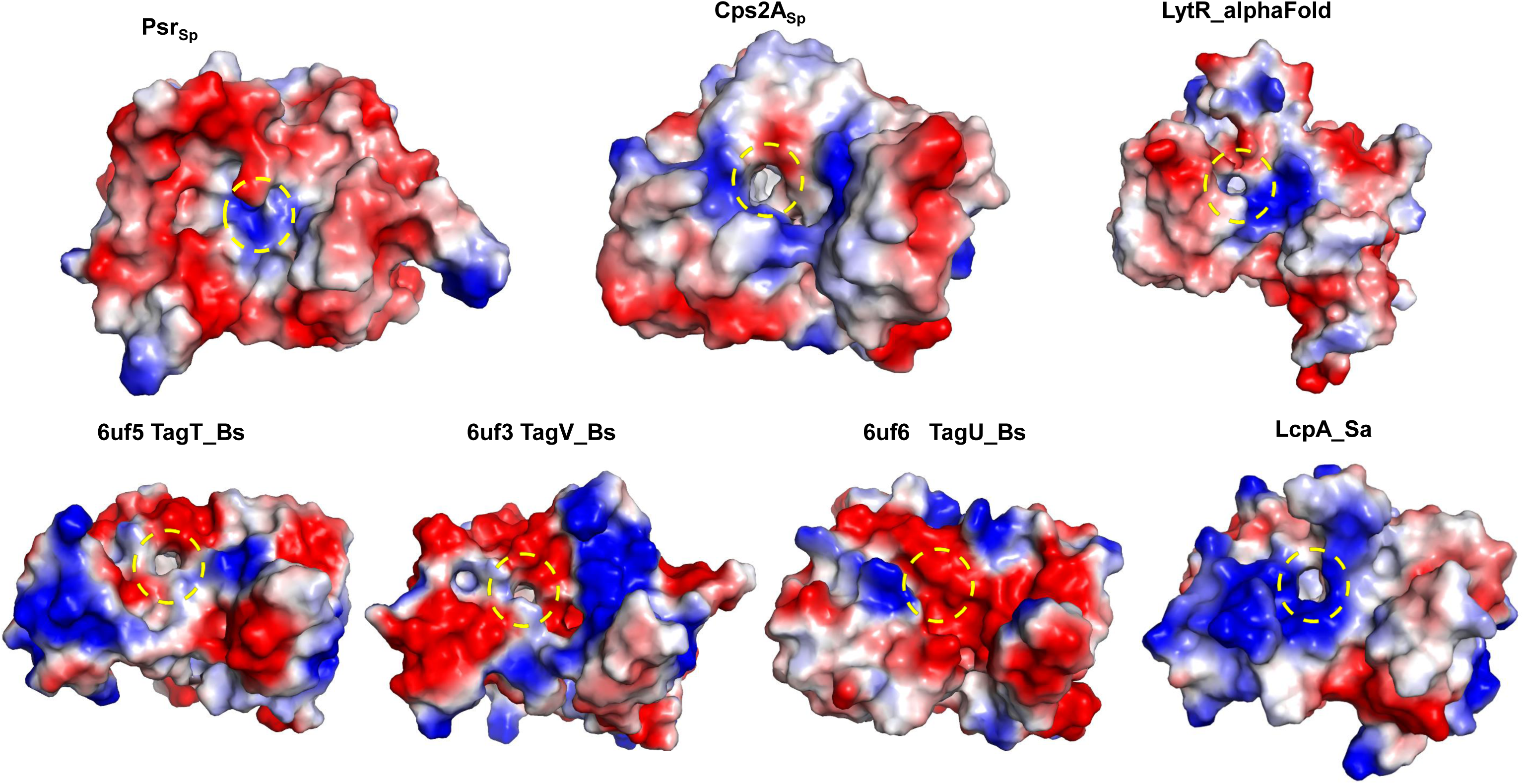
Comparison of electrostatic surface potentials in different LCP indicate different substrate preferences. The surfaces of LCP from different bacterial species, including Psr_Sp_, LcpA_Sa_, Cps2A and TagT_Bs_ are colored according to their electrostatic potential. Positively and negatively charged regions are in blue and red, respectively. The entry of the tunnel for the ligand is indicated by a yellow arrowed circle. This structural comparison indicates differences in sizes at the entry of the tunnel. It also reveals that the entry of the tunnel is surrounded mainly by positively charged residues. Differences in size and in residue charge distribution can be related to different substrate preferences.

It is known that the function of LCP transferases is Mg-dependent (Kawai et al. 2011). The Mg^2+^ ion is not essential for substrate binding, since no ion has been found in some crystal structures of LCP proteins, even if these bound efficiently to the octaprenyl-pyrophosphate moiety of the ligand. However, presence of Mg^2+^ increases significantly phosphotransferase activity (10). The crystal structures of TagT_Bs_ (13) and Cps2A (10) demonstrate that the Mg^2+^ ion binds to six oxygen atoms in perfect tetrahedral symmetry, with two oxygen atoms from the pyrophosphate moiety, two from water molecules and two from the conserved aspartate residues. Comparison of the structures of the same protein with and without Mg^2+^ ion revealed that ion binding modifies the conformation not only of the two aspartates directly involved in the Mg^2+^ binding but also of several arginine residues which bind the pyrophosphate moiety. Thus, Mg^2+^ stabilizes the active conformation of the active site facilitating catalysis. Multiple sequence alignment as well as comparison of the crystal structures of Psr_Sp_ with Csp2A and TagT clearly illustrate that residues D152 and D165, located at the C-terminus of the β1-strand and at the N-terminus of the β2-strand, respectively, are Mg^2+^ binders (**Figure 1**).

The poly-isoprenyl-binding tunnel contains several LCP-conserved residues, including G150, F247, F250, Q329, and V332. Although other residues lining the tunnel, such as M168, L170, M179, L333, I336, L337, F357, V364, L369, V376, and L377 are not conserved among LCP proteins, they keep a hydrophobic nature within the tunnel that is favorable for binding to the undecaprenyl moiety (**Figure 3C**). In addition to the pocket aimed for the hydrophobic carrier bearing the TA, LCPs have a binding site for the second substrate, the PG. Based on comparative analyses of crystal structures of different LCPs, the PG-binding site was proposed to be localized between regions A, C and D (**Figures 3A and 3B**), with the four conserved residues R163, D187, K208 and D322 that could interact with the PG moiety in Psr_Sp_.

### Differential flexibility of the four ABCD regions explains the ability of LCPs to adapt to different TA and PG substrates

The actual substrates of LCP enzymes are much longer than the compounds found in the LCP crystal structures that have been determined until now. To illustrate the relative size of the teichoic acid (TA), the peptidoglycan moiety (PG) and the LCP molecule, a hypothetical molecular model of the Psr_Sp_ complex with these two substrates was created (**Figure S1**). Our model clearly indicates that regions A and B could be very important for TA binding and that region D is positioned close to the PG. Comparison of substrate-free LCPs with different LCP/substrate complexes revealed that the binding of the undecaprenyl-PP compound to LCPs does not require large conformational changes within the protein (10, 12, 13). However, the importance of the mobility of regions A, B, C, and D for binding to the full-length LCP substrates remains unclear. Region A is very flexible in the substrate-free Psr_Sp_, and the conformation of the stretch of residues 153-162 depends on the environment, since it is different in the three molecules found in the asymmetric unit of our Psr_Sp_ crystal (**Figure 2A**). In other ligand-free LCPs such as TagT (6uf5.pdb), region A is not visible in the electron density. High flexibility of the B-region also follows from the fact that residues 288-291 are not visible in the electron density in two out of the three chains of Psr_Sp_ in the asymmetric unit. Poor electron density represents an extreme case of flexibility, and more limited differences in the mobility of the protein within the crystal can be estimated by differences in temperature B-factors (21). The analysis of the B-factor values along the protein chain can identify flexible regions in the protein. **Figure 5** presents the crystal structures of several LCP proteins colored according to their B factors with rigid residues/sections (low B-factor) displayed in blue and flexible regions (higher B-factor) in red. This comparative analysis demonstrates the rigidity of the central core of the Psr_Sp_ protein (comprising the central β-sheet and the helices α1, α2, α8). Regions C and D are rigid in Psr_Sp_, while regions A and B are significantly more flexible. Interestingly, all three molecules within the asymmetric unit of the Psr_Sp_ crystal have a similar distribution of temperature B-factors over their structures. Remarkably, similar analyses of other LCP molecules revealed different rankings of mobile regions (**Figure 5**). Region A is not very flexible compared to region D in Csp2A, whereas region A is the most rigid and region D is the most flexible in LcpA_Sa_. Comparison of four different crystal structures of TagT_Bs_, crystallized both without and with the undecaprenyl-PP-sugar compounds, demonstrated that the presence of these ligands does not alter the range of flexibility of the ABCD regions.

**Figure 5.**
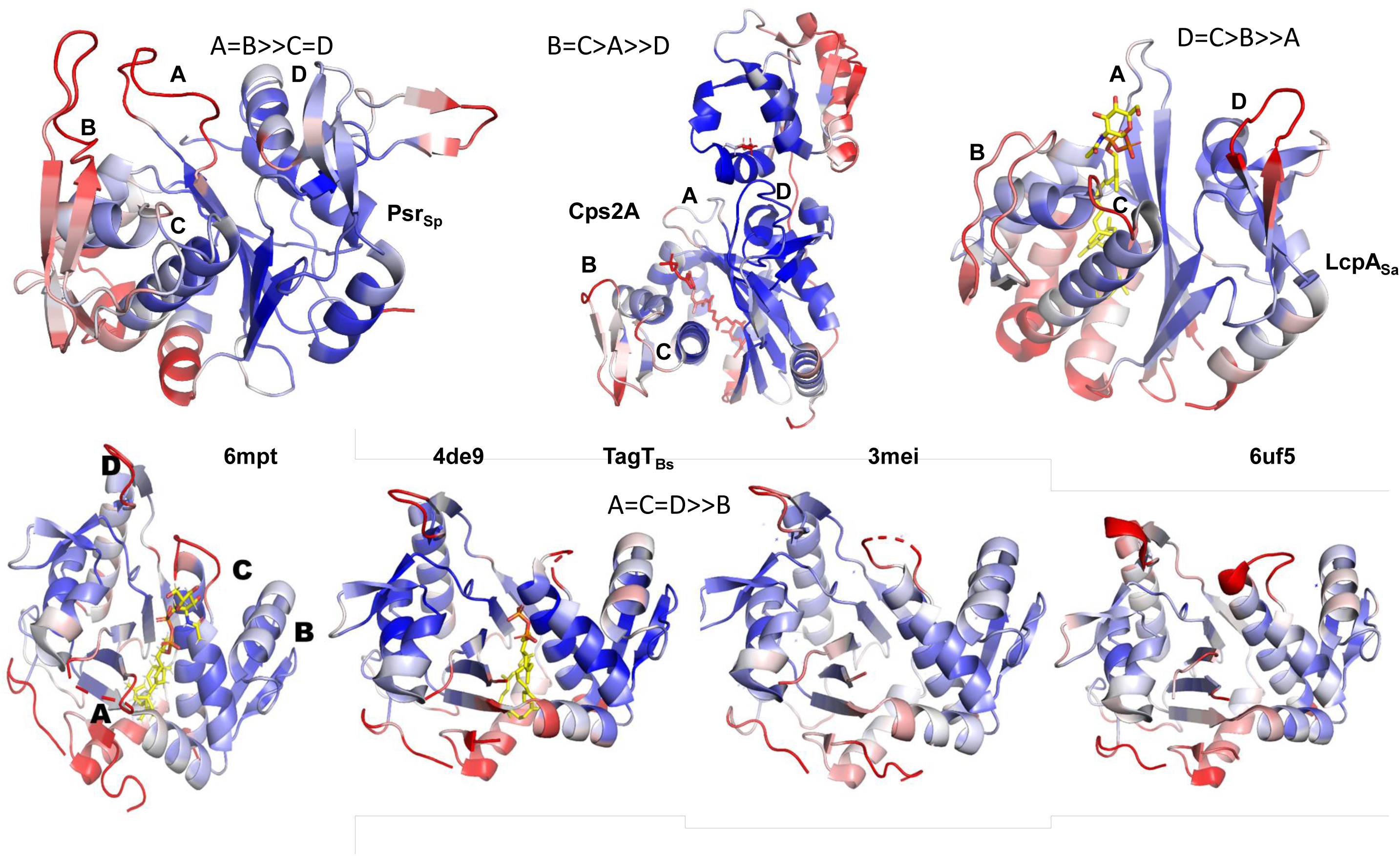
Comparison of crystallographic temperature B-factors in LCP crystal structures demonstrates differences in the mobility of specific loops. The rigid parts of each LCP protein are colored in blue, while the flexible parts are in red. This comparison allowed us to differentiate between the relative flexibility of the loops in the LCP proteins, allowing us to establish patterns. While loops A and B are significantly more flexible than C and D in Psr_Sp_ (A=B>>C=D), loops B and C are more flexible than loop A, which in turn is much more flexible than loop D in Csp2A from *S. pneumoniae* (B=C>A>>D). In contrast, the LCP protein LcpA from *S. Aureus* has a pattern in which D=C>B>>A.

B-factor values characterize the flexibility of a protein in the crystalline state. Nuclear magnetic resonance (NMR) experiments can allow us to assess the mobility of protein domains in solution. Therefore, in an effort to evaluate the dynamics of Psr_Sp_ in solution, we performed NMR experiments to assign the backbone nuclei of this LCP.

### Successful assignment of backbone Psr_Sp_ resonances allows dynamic studies

The validity of the secondary structure analysis based on chemical shifts (CS) depends on the knowledge of the chemical shifts (CS) of the ^1^H(N), ^15^N, ^13^Cα, ^13^Cβ, ^13^C’ nuclei. Our crystal structure revealed that the secondary structure of Psr_Sp_ is composed of a five-stranded β-sheet (β6-β1-β2-β3-β10) (**Figure 2B**), surrounded by eight α-helices and several potentially disordered regions. Folded and intrinsically disordered regions of proteins exhibit different relaxation properties, requiring optimal NMR conditions tailored to each region. This necessitates finding the best sample preparation conditions to achieve the highest quality NMR experiments. Additionally, it is well established that the amide ^1^H and ^15^N chemical shifts in disordered regions and loops are highly sensitive to buffer conditions, including pH and temperature. To optimize data collection, we first performed NMR experiments at two different temperatures, 298K and 308K. We also conducted experiments in a physiological buffer at pH 6.8, with low salt concentration. These conditions are optimal for future analysis of Psr_Sp_ protein interactions with ligands.

NMR data collection showed that the Psr_Sp_ sample remained stable at temperatures up to 313K. No loss of signal intensity in the NMR spectra was detected over a period of two years. A notable issue was the recovery of amide protons in the U-[^15^N,^13^C,^2^H]-labeled samples of Psr_Sp_, which took us nearly a year to achieve full proton recovery in some cases. These amide protons belong mainly to amino acids involved in hydrogen bonds within the β-strand core of the protein, shielding them from water exposure. Several refolding processes were used to potentially overcome this problem, but this approach was ultimately not applicable since Psr_Sp_ did not fully fold into its starting conformation. Nevertheless, we used these U-[^15^N,^13^C,^2^H]-labeled samples of Psr_Sp_ to assign backbone resonances in less crowded, simplified NMR spectra due to the absence of ^2^H-N amide resonances. As a next step, we made use of 75% U-[^15^N,^13^C,^2^H]-labeled samples of Psr_Sp_ to detect and assign the missing ^1^H-^15^N backbone resonances. As a control for the recovery of all ^1^H-^15^N amide cross peaks in the U-[^15^N,^13^C,^2^H]-labeled samples of Psr_Sp_, we additionally analysed the 2D TROSY spectrum of these U-[^15^N,^13^C]-labeled Psr_Sp_ samples.

Since no program is available for automatic assignment of large proteins with both folded and disordered regions, we used a conventional manual assignment strategy based on an approach described in the material and methods section. To achieve the best resolution in the 3D experiments, especially for disordered regions, NMR experiments were mostly performed with the NUS option (22). The ^1^H-^15^N TROSY HSQC spectrum at 308K shows well-dispersed and narrow line widths for the amide signals for the 100% U-[^15^N,^13^C,^2^H]-labeled samples of Psr_Sp_ (**Figure 6**). Under these conditions, with the back-exchange of ^2^H to ^1^H, we observed and assigned 259 out of 295 amino acids, including prolines (**Figure 6D**). 79% ^1^H(N) and ^15^N excluding proline, 87% of ^13^Cα, 93% of ^13^Cβ and 85% of ^13^C’ were assigned successfully. All ^1^H, ^15^N and ^13^C chemical shifts of the LCP domain of Psr_Sp_ at pH 6.8 and at 308K have been deposited in BioMagResBank (http://www.bmrb.wisc.edu) under the accession code BMRB ID 52556. Although the assignment of 36 Psr_Sp_ amino acid residues is still missing and distributed throughout the structure (marked in red in **Figure 6D**), we successfully assigned signals for the residues in the structural regions of interest. Backbone resonances were fully assigned in the B and D regions (**Figure 6D**), while all backbone resonances were also assigned for region A except for the NH groups of residues D152 and S159. Finally, most of the backbone resonances were also assigned for the C region, Additionally, all LCP-conserved residues, which are located close to the polar moiety of the ligand (including D152, R154, R163, D165, S166, R186, D187, K208, R314, R316 and R325), were successfully assigned except for D322. Similarly, most of the Psr_Sp_ hydrophobic residues that surround the hydrophobic part of the ligand (including M168, L170, M179, F247, F250, Q329, L333, I336, L337, and V364) were fully assigned. In contrast, the hydrophobic residues L369, V376, G150, V332, and F357 were not found in the spectra, likely due to incomplete recovery of amide protons.

**Figure 6.**
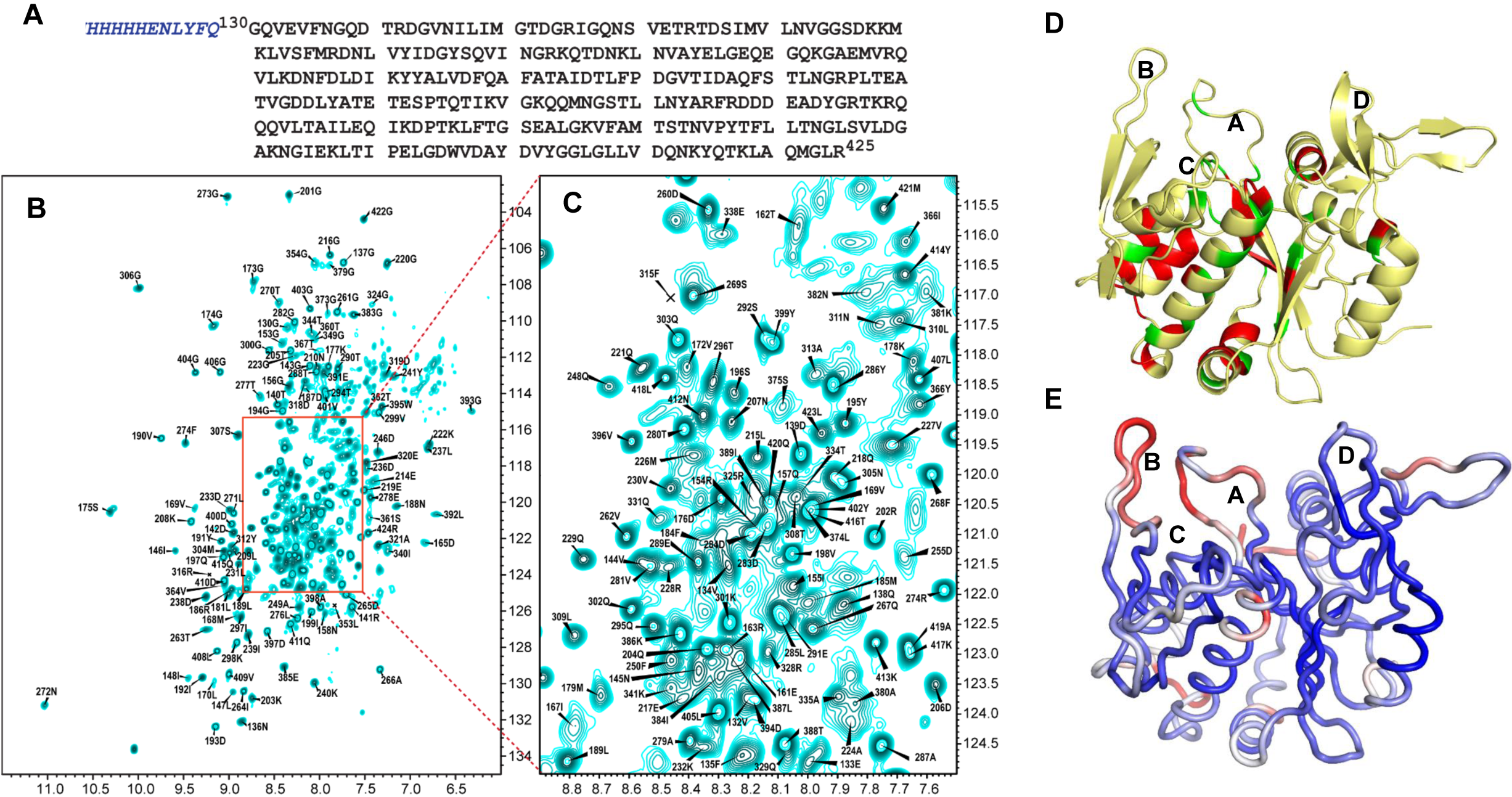
^1^H-^15^N TROSY HSQC spectra of Psr_Sp_ at T=308K with backbone amide assignments. **A.** The amino acid sequence of the Psr_Sp_ construct used in our NMR analyses is presented. Residues from the His-tag are indicated in italic blue. **B.** ^1^H-^15^N TROSY HSQC spectrum at T=308K with its extended crowded part (**C.**) from the ^15^N, ^13^C, ^2^H labelled Psr_Sp_ at 0.7 mM concentration in a 25 mM sodium phosphate buffer (Na^2^HPO ^+^NaH PO) PO ^3-^, pH 6.8 containing 100 mM NaCl, 1 mM NaN_3_, 10 (v/v) % D_2_O and 0.1 mM DSS. The chemical shift assignment of NH backbone is shown by the number and amino acid symbols corresponding to the sequence. The assignment is presented only for residues which cross-peaks are observed at T=308K. **D.** Residues of Psr_Sp_ with fully assigned peaks in the NMR spectrum are displayed in yellow. Carbon-assigned residues are shown in green, and non-assigned residues in red. **E.** Overall structure of Psr_Sp_ colored in red-white-blue according to the predicted order parameter S^2^ with flexible regions in red, and rigid regions in blue.

### Secondary structure of the Psr_Sp_ protein in solution

The chemical shifts of the backbone of the full-length Psr_Sp_ protein were analyzed using the TALOS N program (23) (**Figure S2**). Our results demonstrate that the secondary structure analysis of the folded domain of Psr_Sp_ from our NMR data closely matches the crystal structure of Psr_Sp_, validating our resonance assignments. 12 β-strands and seven α-helices were identified in the NMR-derived secondary structure. Small variations at the beginning and end of the α and β structural elements of Psr_Sp_ can be attributed to environmental differences, including a significant temperature difference of approximately 100K between our NMR measurements and the crystallographic conditions. Notably, the fully unassigned region between residues 242-258, corresponding to the β6 strand and α3 helix in the crystal structure, showed no amide signals in the NMR spectra due to incomplete deuterium exchange. However, TALOS N predicted the secondary structure of this region based only on its sequence, and it matched the expected structure (**Figure S2**). Our attempts to simulate chemical shifts of the Psr_Sp_ protein backbone using the ShiftX2 program (24), and re-perform TALOS N with a combined dataset of experimental and predicted shifts in the region between residues 242-258 yielded similar results (**Figure S2**).

Next, we focused on the structural regions A-D, where amino acid assignments were either fully complete (regions B and D) or nearly complete (regions A and C) (**Figure S2**). Notably, some residues in region B (285–292) could not be observed in the crystal structure of Psr_Sp_, rendering our NMR data the primary source of structural information for these segments. In regions B and D, the short β-strands β7-β9 and β10-β12, were identified in both the crystal structure and the NMR secondary structure (**Figure S2**). Region A is predicted by TALOS-N as a random coil element, which is also consistent with the results from the crystal structure.

To further investigate the conformational flexibility in regions A-D of Psr_Sp_, NMR spectroscopy offers detailed, site-specific insights into protein motions across various time scales. Traditionally, the measure of protein dynamics using NMR involves experimental data like NOEs, T1, and T2, which are interpreted using model-free parameters (25, 26). These parameters describe the spatial and temporal motion of N-H(N) amide bonds in the protein backbone, with order parameters (S²) ranging from 0 to 1. An S² value between 0.8 and 1.0 indicates limited motion, while correlation times describe motions on the ps-ns timescale. A simplified alternative method based on the backbone ^15^N and ^13^C nuclei chemical shifts has been previously proposed (27) and thereafter implemented in TALOS (23). This enables for accurate, site-specific mapping of protein backbone mobility without the need for extensive NMR relaxation experiments beyond standard backbone assignments. This approach, known as the Random Coil Index (RCI) (28), predicts the model-free backbone order parameters S². In the present study, these predicted order parameters are presented in **Figure S2** and mapped on the three-dimensional structure of Psr_Sp_ (**Figure 6E**).

For secondary structural elements like β-strands and α-helices, the predicted order parameters S² are typically high, around 0.8, indicating their rigidity. However, in regions A-D of Psr_Sp_, differences in flexibility are evident. Region D consists of a twisted antiparallel beta sheet formed between the β11-β12 strands, connected by a loop between residues 400-405. The predicted S² values for the amide bonds in both the β sheet and the loop are close to 0.8, indicating low mobility on the ps-ns timescale. This suggests that region D has a stable conformation, which may play a role in substrate recognition and selectivity in the Psr_Sp_ protein. Region C, characterized by a sharp turn between helices α4 and α5, has predicted S² values ranging from 0.7 to 0.8. Although these values are still relatively high, the prediction confidence from TALOS analysis is slightly lower due to incomplete assignment in this region compared to the fully assigned region D. Nonetheless, the rigidity of the C-turn might be crucial for maintaining the correct positioning of α4 and α5 in the active site.

A more complex conformational behaviour is observed in region B, which spans the longest chain of residues (262–304) and has the most complete NMR assignment. Consistent with both X-ray and NMR data, the secondary structure of this region includes two antiparallel β-sheets, formed by the β7-β9a and β8-β9 strands, with higher predicted S² values, indicating rigidity. Region B contains also two coil elements. The first is a loop between residues 286-294 connecting the β8-β9 strands, with low S² values, which suggests high flexibility. This loop is part of the substrate recognition site in the protein, and its conformational plasticity allows it to accommodate different substrates through a conformational selection mechanism. It is however possible that the entire element, including the β8-β9 strands and the loop, experiences slow dynamics, which were not fully explored in this study.

An interesting discrepancy between the X-ray and NMR data is observed in the region comprising residues 267-275. This stretch of residues connecting the β8-β9a strands is disordered in the crystal structure. However, the β7 strand is extended by five amino acids in the NMR structure, with higher S² values, followed by a short turn between residues 271-275, where the S² value decreases to 0.6. The reasons for these differences remain unclear and cannot be fully explained by the data obtained in this study, suggesting the need for further investigation. Region A which corresponds to a loop that connects the β1 and β2 strands, does not contain any secondary structure elements, as confirmed by both X-ray and NMR data. The predicted S² values drop to as low as 0.4, indicating its conformational heterogeneity and plasticity. We suggest that the close spatial proximity of the flexible loop in region A to the similarly flexible loop in region B is key to the capacity of Psr_Sp_ to accommodate various substrate shapes within the active site.

## Discussion

LCP proteins are attractive therapeutic targets since these enzymes catalyze a reaction essential for Gram-positive bacteria, resulting in the attachment of a secondary cell wall glycopolymer to the peptidoglycan. Inhibition of this process disrupts cell wall assembly, cell division and pathogenicity. Importantly, since LCP enzymes are located on the extracellular part of bacterial cell, inhibitors of LCP enzymes are not required to cross the membrane which renders the creation of antimicrobial compounds significantly easier. Here, we describe the crystal structure and NMR dynamic analyses of Psr_Sp_, the second LCP enzyme from *S. pneumoniae* that catalyzes attachment of wall teichoic acid as well as capsular polysaccharides to the PG. The crystal structure of Psr_Sp_ reveals a typical LCP fold, despite low sequence homology. Comparison with several other LCP structures confirmed that substrate binding and functional mechanisms are the same as in all other LCP proteins. The conservation of the residues participating in the catalysis suggests a common mechanism for this reaction. Thus, all the hitherto gathered structural and mutational data from different LCP reveal a common binding mode of the pyrophosphate-lipid carrier moiety by LCP transferases. However, it should be noted that the real substrates of LCP enzymes are much larger than the ligands found in crystal structures. Moreover, their chemical composition is different not only for different bacteria but even within the same organism. For example, there are almost 100 serotypes described for *S. pneumoniae*, with different capsular polysaccharides on their surfaces (29). Many of these are covalently linked to the PG in the same way as for WTA. Though it has still not been demonstrated experimentally that LCP transferases catalyze the reaction for WTA and all capsular polysaccharides, LCP genetic and bioinformatics suggest that it is true.

Most of the bacteria contain three genes coding for three different LCP enzymes. Differences in the electrostatic potential on the surface of the three LCP from *S. pneumoniae* (**Figure 4**) could reflect the different specificity of each LCP. It has been shown that all three LCP from *S. aureus* can catalyze the attachment of the staphylococcal WTA to the PG (8). However, LcpA is the most important enzyme among these three LCPs for the attachment of WTA (13). Interestingly, another LCP enzyme, LcpC, is the most important for the PG attachment of the type 5 capsular polysaccharide in this bacterium (30). Similarly, all three LCP proteins contribute to the catalysis of the reaction between PG and secondary glycopolymers in *S. pneumoniae* (9). However, we still do not know the variation nor the basis in the preference of the catalysis performed by the different LCP for different polysaccharides.

We hypothesize here that the mobility of the loops interacting with the polysaccharide could be of the greatest importance to create a promiscuous binding. The connection between flexibility of proteins and promiscuity of their interactions is well-established for intrinsically disordered proteins (31, 32). Interactions via flexible regions of separate molecules which become ordered during assembly allows the same protein to interact with multiple partners. A similar relation was found for several enzymes when increased flexibility of their active sites resulted in increased numbers of possible substrates (33, 34). Two methods were used in the present study to estimate the mobility of the essential regions of Psr_Sp_. First, flexibility was estimated using B-factor values obtained for the crystal structure of Psr_Sp_. Second, flexibility was also estimated for Psr_Sp_ in solution from predicted order parameter S^2^ obtained from chemical shifts in the NMR spectra.

Consistency in the secondary structure elements obtained from the crystal structure and NMR experiments despite a temperature difference of 200°C between the conditions used in the different experimental procedures, evidence for the stability of the Psr_Sp_ molecule and for the correct assignment of the peaks in our ^1^H-^15^N TROSY HSQC spectrum. In this study we found that the chemical shifts predicted by ShiftX2 for rigid secondary structures in proteins closely match those obtained through NMR, particularly for the ^13^Cα, ^13^Cβ, and ^13^C’ nuclei. This finding suggests that it may not always be necessary to aim for a full assignment of all parts of very large proteins. In large protein production, deuteration is essential but often problematic because fully recovering the amide protons is difficult, expensive, and can lead to protein loss. Since amide recovery mainly affects the most stable parts of the protein, the results from the present study indicate that it might be more efficient for large proteins to focus on recovering amide protons only from flexible regions. This approach could simplify the analysis by reducing spectral crowding.

Our results reveal that loops A and B in Psr_Sp_, responsible for interactions with WTA or capsular polysaccharides, are significantly more mobile compared to loop D which is suggested to be involved in the interaction with the PG. This is not true for the other LCP protein Cps2A from *S. pneumoniae*. Analysis of B-factors in the crystal structure of Cps2A shows that this LCP has a very rigid region A (**Figure 5**) and a short flexible B region, which contrast significantly with Psr_Sp_. Furthermore, the staphylococcal protein LcpA_Sa_ has a rigid region A, a B region with some flexibility, and highly mobile C and D regions (**Figure 5**). We find it interesting that the number of serotypes and the chemical variety of capsular polysaccharides is much smaller in *S. aureus* compared to *S. pneumonia* (35, 36). One of the many remaining questions is thus, how can this flexibility be converted to substrate specificity?

Unfortunately, we do not have enough biological data on the substrate specificity nor on the promiscuity of the three different LCP phosphotransferases. It has been previously shown that all three LCP molecules in *S. pneumoniae* contribute to the maintenance of the normal volume of polysaccharides on the cell surface (9). Deletion of Cps2A is not defective in capsule attachment (37). While a Cps2A mutant of the strain D39 of *S. pneumoniae* produces only 30-40% of the capsule, a Psr_Sp_ mutant produces about 70% of the capsule (9). D39 has a serotype 2 capsule, with its repeating units formed by glucose, glucuronic acid and rhamnose. Although the previous results of Eberhardt *et al* demonstrate that Cps2A is more active with a serotype 2 capsule compared to Psr_Sp_, it still does not provide a definitive answer in regards to which protein is active with more different serotypes.

In conclusion, the present study based on a combination of X-ray crystallography and NMR analyses, revealed that regions A and B in Psr_Sp_ are significantly more flexible compared to other parts of this key LCP protein. We speculate that the flexibility of the A and B loops could be important, providing the LCP Psr_Sp_ with the possibility to catalyze PG attachment for a large array of different glycopolymers. Identification of the most promiscuous LCP enzyme could be of great importance for the development of future antibiotics.

## Experimental Procedures

### Cloning, expression and purification of Psr_sp130-424_

The sequence of the extracellular LCP domain of Psr_Sp_ (residues 130-424; UNIPROT Q8DPD6_STRR6) was cloned into the vector pET28 in frame with a N-terminal His-tag for NMR studies and in frame with a C-terminal His-tag for crystallization assays. Plasmids were transformed into the BL21 (DE3) pLys *E. Coli* strain, and cells were cultured at 37°C in LB broth supplemented with 50 μg/ml kanamycin until OD_600_ of 0.8 was reached. Protein transcription was then initiated by reaching a final concentration of 0.5 mM isopropyl-β-D-1-thiogalactopyranoside (IPTG), which was added to the culture after reducing the culture temperature to 20°C before overnight incubation. Cells were harvested by centrifugation and thereafter resuspended in lysis buffer (20 mM Tris pH 7.5, 250 mM NaCl, 20 mM Imidazole) supplemented with Complete protease inhibitor (Roche). Cells were lysed by sonication, and cell debris were pelleted by centrifugation. The supernatant was loaded on a 5 ml HisTrap FF column (Cytiva) and after column wash, the protein construct Psr_Sp131-424_ was eluted with 500 mM Imidazole and concentrated down to 5 ml with an Amicon Ultra centrifugal filter. The protein was further purified via size exclusion chromatography with a HiLoad 16/60 Superdex, 75 pg and eluted in 20 mM Tris pH 7.5, 150 mM NaCl as a monomeric peak. Peak fractions were pooled and concentrated to 25 mg/ml for crystallization assays. Protein purity was assessed using SDS-PAGE. For NMR studies, the buffer of Psr_Sp131-424_ was exchanged into 25 mM NaPhosphate, 100mM NaCl, and concentrated to 200 μM.

### Crystallization and structure determination

Psr_Sp_ crystals were obtained at 20°C using the sitting drop technique, mixing equal amounts of the protein Psr_Sp_ and the reservoir solution (25 mM NaPhosphate pH 6.8, 100 mM NaCl), in a 400 nl drop. Crystals were cryo-protected by adding 20% ethylene glycol to the crystallization drop and thereafter flash-frozen in liquid nitrogen. Diffraction data were collected at the BioMax Beamline at the MaxIV Laboratory (Lund, Sweden). The crystal diffraction was anisotropic and the integrated data were shared with the STARANISO web server (http://staraniso.globalphasing.org/cgi-bin/staraniso.cgi) for elliptical truncation after data processing in XDSAPP3 to 2.15Å. We were able to determine the crystal structure of Psr_Sp131-424_ by molecular replacement only by using a three-dimensional molecular model generated with AlphaFold (16) as a search template. Three molecules were present in the asymmetric unit of the crystal according to the calculated Matthews coefficient. The first two monomers were found with PHASER (15), while the third monomer was found by MOLREP (17), based on the PHASER output. Model building and refinement were performed with Phenix (Version 1.19.1-4122) (38), CCP4 (14) and Coot (Version 0.9) (39).

### Expression of isotope-labelled Psr_Sp_ and preparation of NMR samples

The DNA sequence encoding for Psr_Sp130–420_ and a N-terminal MHis_6_ENLYFQ-tag was cloned into the pET28a expression vector (Novagen). The tag-Psr_Sp_ construct was transformed into the BL21 (DE3) pLys *E. coli* express competent cells, and the protein was thereafter expressed in different isotopic labelling combinations in ^1/2^H, ^15^N, ^12/13^C-labelled M9 medium. Chemicals for isotope labelling including ammonium chloride, ^15^N (99%), D-glucose, ^13^C (99%) and deuterium oxide were purchased from Cambridge Isotope Laboratories. Psr_Sp_ for NMR experiments was expressed in 1 L D_2_O M9 medium using 3 g/L of U-[^13^C,^2^H]-glucose (CIL, Andover, MA) as the main carbon source, and 1 g/L of ^15^NH_4_Cl (CIL, Andover, MA) as the nitrogen source. Bacterial growth was continued for 16 h at 16°C, and cells were harvested by centrifugation. Cells were resuspended in lysis buffer (20 mM Tris pH 7.5, 250 mM NaCl, 20 mM Imidazole) supplemented with complete protease inhibitor (Roche) and lysed using an ultrasonicator, followed by centrifugation at 40,000 g for 30 min to remove cell debris. The purification steps were the same as for the construct used for the crystallographic study. The final Psr_sp131-424_ protein sample was subsequently exchanged to a buffer composed of 25 mM sodium phosphate (Na_2_HPO_4_ ^+^NaH_2_PO_4_) PO_4_ ^3-^, 100 mM NaCl, pH 6.8, 1 mM NaN_3_, 10 (v/v) % D_2_O, suitable for NMR experiments using gravity flow PD10 desalting columns (GE Healthcare). All NMR experiments were performed by adding 0.1 mM DSS (4,4-dimethyl-4-silapentane-1-sulfonic acid) as an internal ^1^H chemical shift standard. Protein concentration was about 0.7 mM, and spectra were acquired in a 3 mm tube. ^13^C and ^15^N chemical shifts were referenced indirectly to the ^1^H standard using a conversion factor derived from the ratio of NMR frequencies (40).

### NMR experiments

NMR experiments on U-[^15^N,^13^C,^2^H], 75%U-[^15^N,^13^C,^2^H] and U-[^15^N,^13^C] labelled samples of Psr_sp131-424_ were acquired either on a 700 MHz Bruker Avance III spectrometer equipped with a 3 mm cryo-enhanced QCI-F probe, or on a 600MHz Bruker Avance III spectrometer equipped with a 5mm cryo-enhanced QCI-P probe. The experiments were performed at 308K to improve relaxation parameters. Backbone resonance assignments for Psr_sp131-424_ were obtained as previously described (41–43) and were based on a set of 3D TROSY or HSQC triple resonance experiments from the Bruker library. In summary, to increase resolution in the indirect dimensions, and reduce acquisition time for the 3D experiments, the iterative non-uniform sampling protocol (NUS) (22) was used in the experiments comprising TROSY-HNCO, TROSY-HNCA and TROSY-HN(CO)CA, TROSY-HN(CA)CO, TROSY-HN(CO)CACB, TROSY HNCACB, H(CC)(CO)NH, TOCSY-^15^N HSQC, HCACO and TOCSY-^13^C HSQC experiments. Additionally, NOESY ^15^N-HSQC, NOESY-^13^C-HSQC spectra were collected (44–46). The combined 3D NUS NMR target acquisition (TA) data were processed using the IST algorithm in the NUS module in TopSpin4.0.6 (Bruker, Billerica, MA, USA), and analysed using CcpNmr2.4.2.(47) and Dynamics Center2.8 (Bruker, Billerica, MA, USA). Data submitted to the BioMagResBank with accession code BMRB ID is 52556. The chemical shifts (CS) of ^1^H(N), ^15^N, ^13^Cα, ^13^Cβ and ^13^C’ nuclei for every amino acid of Psr_sp131-424_ were analysed with the TALOS-N software (23) to extract the secondary structure prediction (S.S. Prediction) index, random coil index (RCI) (28) order parameter S2, and confidence index. In case of no chemical shift, TALOS-N uses a database of sequences to predict the secondary structure. Standard nomenclature for amino acids of the carbon atoms was used in both the main text and the figures, where ^13^Cα is the carbon next to the carbonyl group ^13^C’ and ^13^Cβ is the carbon next to ^13^Cα (48). Bruker standard pseudo 2D pulse sequence tractf3gpphwg was used to obtain a quick estimate of the rotational correlation time of the Psr_sp_ (19).

## Abbreviations

AlphaFold: an artificial intelligence system that can predict three-dimensional structures of proteins from amino acid sequences
PG: peptidoglycan
LCP: LytR-CpsA-Psr
TA: teichoic acid
LTA: lipoteichoic acid
WTA: wall teichoic acid
CP: capsular polysaccharide
C55-PP: undecaprenyl pyrophosphate
PDB: Protein Data Bank
TM: transmembrane
RMSD: root-mean-square deviation
AATGal: 2-acetamido-4-amino-2,4,6-trideoxy-D-galactose
RU: repeating unit
NMR: nuclear magnetic resonance
TROSY: transverse relaxation optimized spectroscopy
HSQC: heteronuclear single quantum coherence.

## Data availability

The crystal structure coordinates and structural factors for Psr_Sp_ have been deposited to the Protein Data Bank under accession code 9GN3. Assignment data was submitted to the BioMagResBank with accession code BMRB ID is 52556. All other information and data are available from the authors upon request.

## Supporting Information

This article contains supporting information.

## Acknowledgements

We gratefully acknowledge access to the synchrotron beam line BioMax at MAXIV (Lund, Sweden) and to the crystallization facility within the Protein Science Facility, Karolinska Institutet, (http://ki.se.proxy.kib.ki.se/en/mbb/protein-science-facility). The authors thank also the Swedish NMR Centre for access to instruments and technical support.

## Author contributions

BMS produced and isolated all necessary protein variants for both the X-ray and the NMR assays. BMS developed a construct of labelled Psr_Sp_ proteins adequate for NMR analyses. BMS has performed all crystallization assays, collected data and performed the initial processing of the data. MM and TS identified the molecular replacement solutions and performed the refinement of the crystal structure. TS performed all structural analyses. TS, TA and AA have written the original manuscript draft. HGL, EA and BHN contributed to the work on the manuscript. PA and DL contributed with all NMR measurements. TA performed NMR assignments of Psr_Sp_. PA obtained the secondary structure of Psr_Sp_ based on the TALOS analysis. TS and AA conceptualized the project, and supervised different parts of the project.

## Funding and additional information

This work was supported by the Swedish Foundation for Strategic Research grant ITM17-0218 to TA and PA, grant RSF 23-44-10021 to D.M.L, Swedish Cancer Society (21 1605 Pj 01 H), Cancer och Allergi Fonden (10399), and Swedish Research Council (2021-05061 and 2018-02874) to AA

---

**Figure S1.**
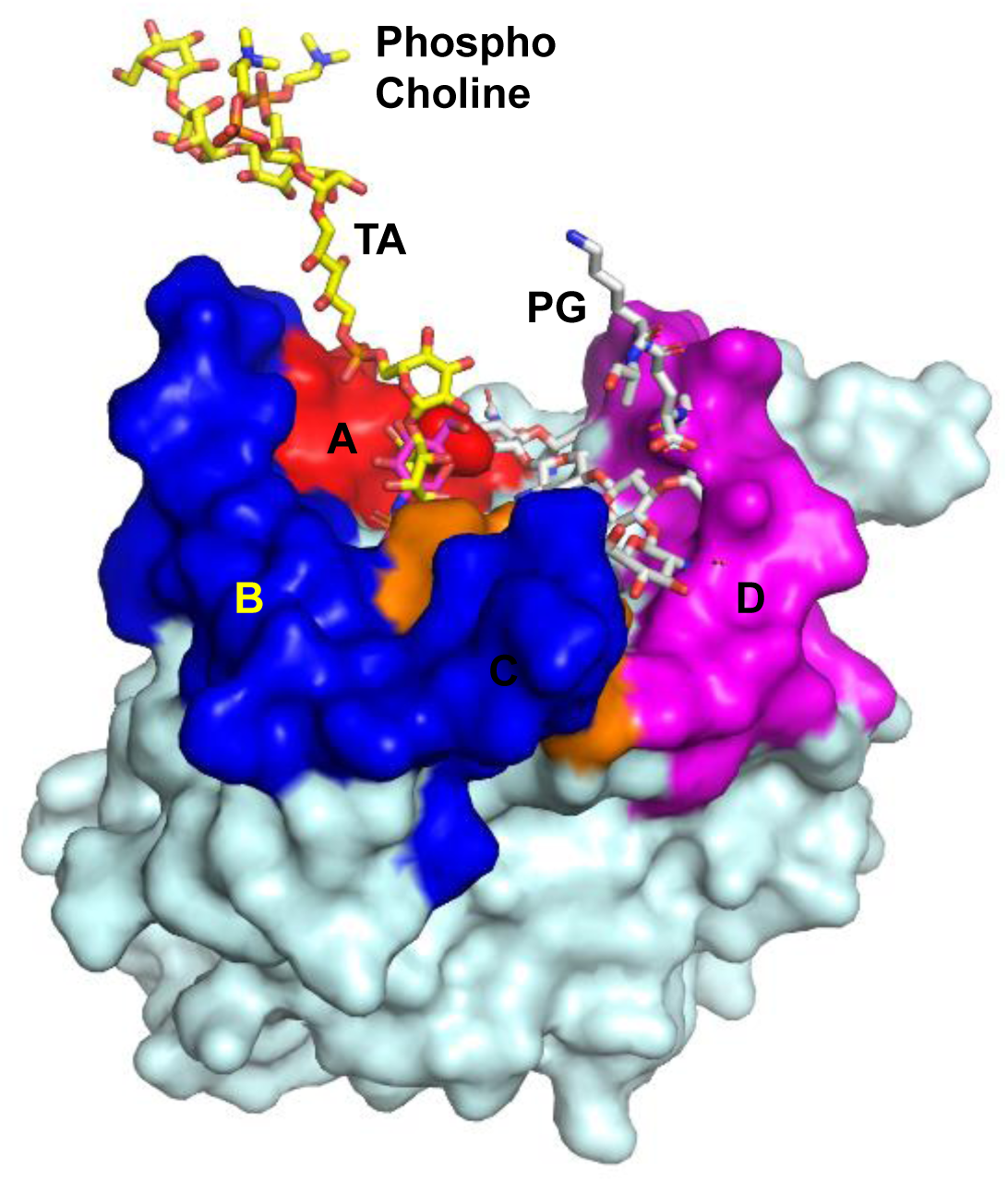
Hypothetical molecular model of the undecaprenol-WTA/PG/Psr_Sp_ complex. demonstrates that regions A and B can interact with large sections of the first RU of polysaccharide whereas region D is close to the peptidoglycan.

**Figure S2.**
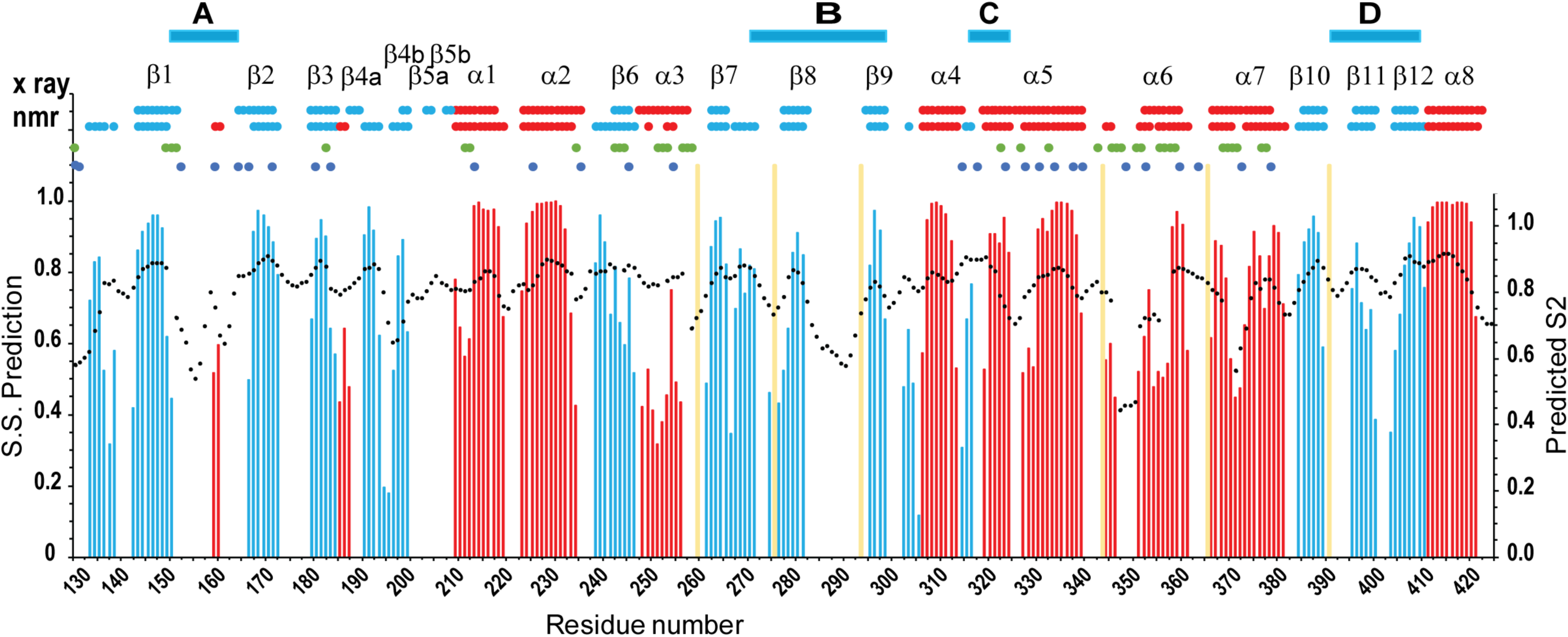
The secondary structures of Psr_Sp_ obtained from NMR is in agreement with those derived from the crystal structure of Psr_Sp_. The secondary structure prediction (S.S. Prediction) index, predicted order parameter S^2^ were extracted from the CS of the ^1^H(N), ^15^N, ^13^Cα, ^13^Cβ and ^13^C’ backbone resonances by TALOS-N. The S.S. Prediction indexes versus the amino acid sequences are presented on the left side of panel. Red and blue bars indicate α-helices/turns and β-strands, respectively. Predicted S^2^ are presented on the right side of the panels, and are indicated by black dots. Schematic representations of the X-ray- and NMR-determined secondary structures are shown on the top of the panels. Blue and green circles represent the amino acids where either only NH or all resonances were not assigned, respectively. The range of the loop regions A, B, C and D are shown on the top of the panels.

**Table S1.**
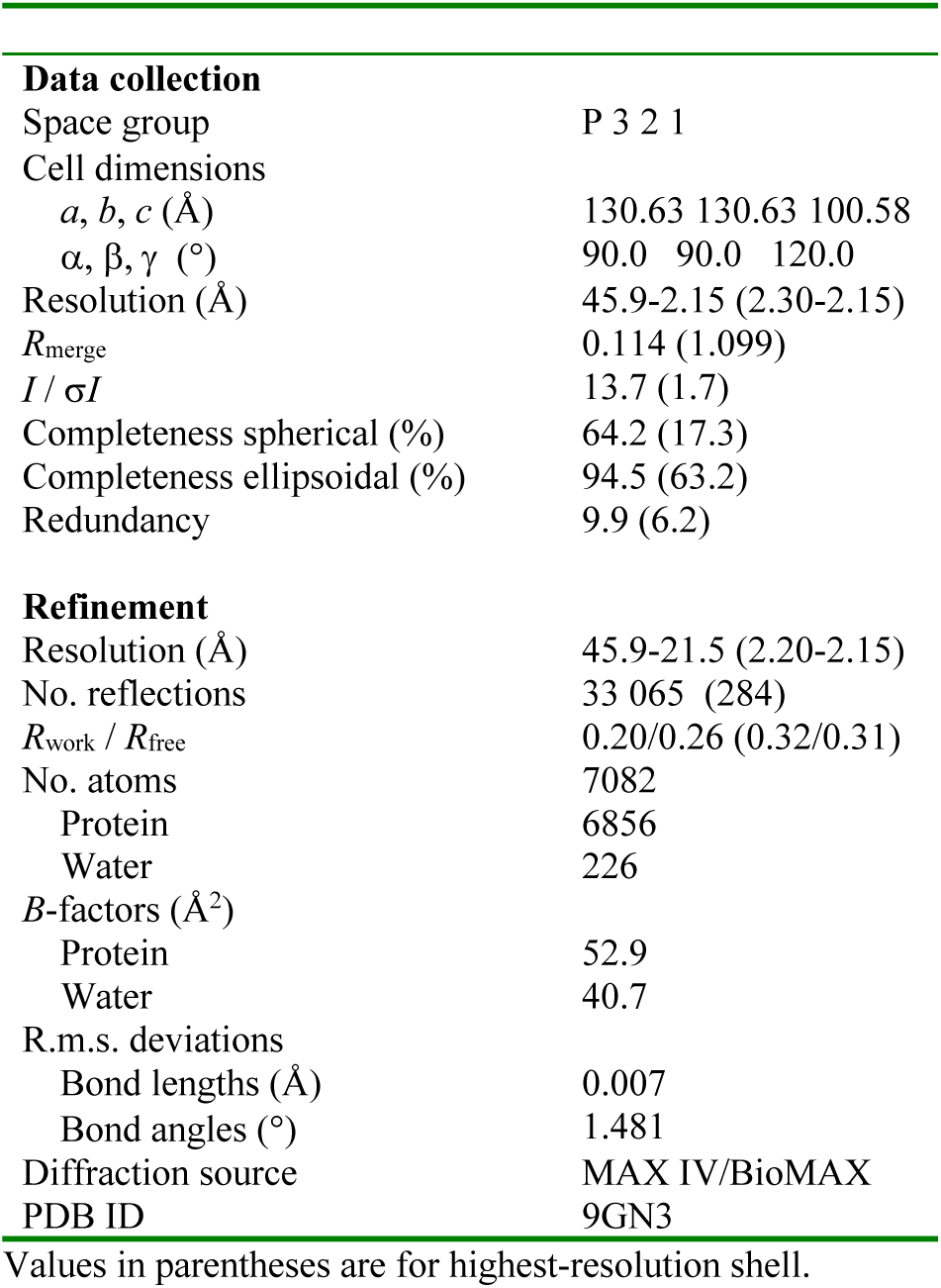
Data collection and refinement statistics for Psr_Sp_.

## References

1. Rajagopal, M., and Walker, S. (2017) Envelope Structures of Gram-Positive Bacteria. Curr Top Microbiol Immunol 404, 1–44

2. Schade, J., and Weidenmaier, C. (2016) Cell wall glycopolymers of Firmicutes and their role as nonprotein adhesins. FEBS Lett 590, 3758–3771

3. Stefanović, C., Hager, F. F., and Schäffer, C. (2021) LytR-CpsA-Psr Glycopolymer Transferases: Essential Bricks in Gram-Positive Bacterial Cell Wall Assembly. Int J Mol Sci 22,

4. Weidenmaier, C., and Peschel, A. (2008) Teichoic acids and related cell-wall glycopolymers in Gram-positive physiology and host interactions. Nat Rev Microbiol 6, 276–287

5. Vollmer, W., Massidda, O., and Tomasz, A. (2019) The Cell Wall of. Microbiol Spectr 7,

6. Aceil, J., and Avci, F. Y. (2022) Pneumococcal Surface Proteins as Virulence Factors, Immunogens, and Conserved Vaccine Targets. Front Cell Infect Microbiol 12, 832254

7. Denapaite, D., Brückner, R., Hakenbeck, R., and Vollmer, W. (2012) Biosynthesis of teichoic acids in Streptococcus pneumoniae and closely related species: lessons from genomes. Microb Drug Resist 18, 344–358

8. Schaefer, K., Matano, L. M., Qiao, Y., Kahne, D., and Walker, S. (2017) In vitro reconstitution demonstrates the cell wall ligase activity of LCP proteins. Nat Chem Biol 13, 396–401

9. Eberhardt, A., Hoyland, C. N., Vollmer, D., Bisle, S., Cleverley, R. M., Johnsborg, O. et al. (2012) Attachment of capsular polysaccharide to the cell wall in Streptococcus pneumoniae. Microb Drug Resist 18, 240–255

10. Kawai, Y., Marles-Wright, J., Cleverley, R. M., Emmins, R., Ishikawa, S., Kuwano, M. et al. (2011) A widespread family of bacterial cell wall assembly proteins. EMBO J 30, 4931–4941

11. Rajaei, A., Rowe, H. M., and Neely, M. N. (2022) The LCP Family Protein, Psr, Is Required for Cell Wall Integrity and Virulence in. Microorganisms 10,

12. Li, F. K. K., Rosell, F. I., Gale, R. T., Simorre, J. P., Brown, E. D., and Strynadka, N. C. J. (2020) Crystallographic analysis of. J Biol Chem 295, 2629–2639

13. Schaefer, K., Owens, T. W., Kahne, D., and Walker, S. (2018) Substrate Preferences Establish the Order of Cell Wall Assembly in Staphylococcus aureus. J Am Chem Soc 140, 2442–2445

14. Agirre, J., Atanasova, M., Bagdonas, H., Ballard, C. B., Baslé, A., Beilsten-Edmands, J. et al. (2023) The CCP4 suite: integrative software for macromolecular crystallography. Acta Crystallogr D Struct Biol 79, 449–461

15. McCoy, A. J., Grosse-Kunstleve, R. W., Adams, P. D., Winn, M. D., Storoni, L. C., and Read, R. J. (2007) Phaser crystallographic software. J Appl Crystallogr 40, 658–674

16. Jumper, J., Evans, R., Pritzel, A., Green, T., Figurnov, M., Ronneberger, O. et al. (2021) Highly accurate protein structure prediction with AlphaFold. Nature 596, 583–589

17. Vagin, A., and Teplyakov, A. (2010) Molecular replacement with MOLREP. Acta Crystallogr D Biol Crystallogr 66, 22–25

18. Cavanagh, J., Fairbrother, W., Palmer III, A., Rance, M., and Skelton, N. (2007) Principles and Practice: Protein NMR Spectroscopy Elsevier Academic Press,

19. Lee, D., Hilty, C., Wider, G., and Wüthrich, K. (2006) Effective rotational correlation times of proteins from NMR relaxation interference. J Magn Reson 178, 72–76

20. Holm, L., Laiho, A., Törönen, P., and Salgado, M. (2023) DALI shines a light on remote homologs: One hundred discoveries. Protein Sci 32, e4519

21. Sun, Z., Liu, Q., Qu, G., Feng, Y., and Reetz, M. T. (2019) Utility of B-Factors in Protein Science: Interpreting Rigidity, Flexibility, and Internal Motion and Engineering Thermostability. Chem Rev 119, 1626–1665

22. Orekhov, V., and Jaravine, V. A. (2011) Analysis of non-uniformly sampled spectra with multi-dimensional decomposition. Progress in Nuclear Magnetic Resonance Spectroscopy 59, 271–292

23. Shen, Y., and Bax, A. (2013) Protein backbone and sidechain torsion angles predicted from NMR chemical shifts using artificial neural networks. Journal of Biomolecular Nmr 56, 227–241

24. Han, B., Liu, Y. F., Ginzinger, S. W., and Wishart, D. S. (2011) SHIFTX2: significantly improved protein chemical shift prediction. Journal of Biomolecular Nmr 50, 43–57

25. Lipari, G., and Szabo, A. (1982) Model-Free Approach to the Interpretation of Nuclear Magnetic-Resonance Relaxation in Macromolecules .2. Analysis of Experimental Results. J Am Chem Soc 104, 4559–4570

26. Lipari, G., and Szabo, A. (1982) Model-Free Approach to the Interpretation of Nuclear Magnetic-Resonance Relaxation in Macromolecules .1. Theory and Range of Validity. J Am Chem Soc 104, 4546–4559

27. Berjanskii, M. V., and Wishart, D. S. (2005) A simple method to predict protein flexibility using secondary chemical shifts. J Am Chem Soc 127, 14970–14971

28. Berjanskii, M. V., and Wishart, D. S. (2005) A simple method to predict protein flexibility using secondary chemical shifts. J Am Chem Soc 127, 14970–14971

29. Paton, J. C., and Trappetti, C. (2019) Capsular Polysaccharide. Microbiol Spectr 7,

30. Chan, Y. G., Kim, H. K., Schneewind, O., and Missiakas, D. (2014) The capsular polysaccharide of Staphylococcus aureus is attached to peptidoglycan by the LytR-CpsA-Psr (LCP) family of enzymes. J Biol Chem 289, 15680–15690

31. Tompa, P., Schad, E., Tantos, A., and Kalmar, L. (2015) Intrinsically disordered proteins: emerging interaction specialists. Curr Opin Struct Biol 35, 49–59

32. Piovesan, D., Monzon, A. M., Quaglia, F., and Tosatto, S. C. E. (2022) Databases for intrinsically disordered proteins. Acta Crystallogr D Struct Biol 78, 144–151

33. McAuley, M., Kristiansson, H., Huang, M., Pey, A. L., and Timson, D. J. (2016) Galactokinase promiscuity: a question of flexibility?. Biochem Soc Trans 44, 116–122

34. Gatti-Lafranconi, P., and Hollfelder, F. (2013) Flexibility and reactivity in promiscuous enzymes. Chembiochem 14, 285–292

35. O’Riordan, K., and Lee, J. C. (2004) Staphylococcus aureus capsular polysaccharides. Clin Microbiol Rev 17, 218–234

36. Visansirikul, S., Kolodziej, S. A., and Demchenko, A. V. (2020) Staphylococcus aureus capsular polysaccharides: a structural and synthetic perspective. Org Biomol Chem 18, 783–798

37. Morona, J. K., Morona, R., and Paton, J. C. (2006) Attachment of capsular polysaccharide to the cell wall of Streptococcus pneumoniae type 2 is required for invasive disease. Proc Natl Acad Sci U S A 103, 8505–8510

38. Adams, P. D., Afonine, P. V., Bunkóczi, G., Chen, V. B., Davis, I. W., Echols, N. et al. (2010) PHENIX: a comprehensive Python-based system for macromolecular structure solution. Acta Crystallogr D Biol Crystallogr 66, 213–221

39. Emsley, P., Lohkamp, B., Scott, W. G., and Cowtan, K. (2010) Features and development of Coot.. Acta Crystallogr D Biol Crystallogr 66, 486–501

40. Wishart, D. S., Bigam, C. G., Yao, J., Abildgaard, F., Dyson, H. J., Oldfield, E. et al. (1995) 1H, 13C and 15N chemical shift referencing in biomolecular NMR. J Biomol NMR 6, 135–140

41. Agback, P., Lesovoy, D. M., Han, X., Sun, R. H., Sandalova, T., Agback, T. et al. (2022) H-1, C-13 and N-15 resonance assignment of backbone and IVL-methyl side chain of the S135A mutant NS3pro/NS2B protein of Dengue II virus reveals unique secondary structure features in solution. Biomolecular Nmr Assignments 16, 135–145

42. Unnerstale, S., Nowakowski, M., Baraznenok, V., Stenberg, G., Lindberg, J., Mayzel, M. et al. (2016) Backbone Assignment of the MALT1 Paracaspase by Solution NMR. Plos One 11,

43. Agback, T., Dominguez, F., Frolov, I., Frolova, E. I., and Agback, P. (2021) 1H, 13C and 15N resonance assignment of the SARS-CoV-2 full-length nsp1 protein and its mutants reveals its unique secondary structure features in solution. PLoS One 16, e0251834

44. Kay, L. E., Ikura, M., Tschudin, R., and Bax, A. (1990) 3-Dimensional Triple-Resonance Nmr-Spectroscopy of Isotopically Enriched Proteins. J Magn Reson 89, 496–514

45. Kay, L. E., Keifer, P., and Saarinen, T. (1992) Pure Absorption Gradient Enhanced Heteronuclear Single Quantum Correlation Spectroscopy with Improved Sensitivity. J Am Chem Soc 114, 10663–10665

46. Schleucher, J., Schwendinger, M., Sattler, M., Schmidt, P., Schedletzky, O., Glaser, S. J. et al. (1994) A general enhancement scheme in heteronuclear multidimensional NMR employing pulsed field gradients. J Biomol NMR 4, 301–306

47. Vranken, W. F., Boucher, W., Stevens, T. J., Fogh, R. H., Pajon, A., Llinas, M. et al. (2005) The CCPN data model for NMR spectroscopy: development of a software pipeline. Proteins 59, 687–696

48. Markley, J. L., Bax, A., Arata, Y., Hilbers, C. W., Kaptein, R., Sykes, B. D. et al. (1998) Recommendations for the presentation of NMR structures of proteins and nucleic acids--IUPAC-IUBMB-IUPAB Inter-Union Task Group on the standardization of data bases of protein and nucleic acid structures determined by NMR spectroscopy. Eur J Biochem 256, 1–15

